# Distinct genetic architectures underlie divergent thorax, leg, and wing pigmentation between *Drosophila elegans* and *D. gunungcola*

**DOI:** 10.1101/2020.06.28.176735

**Authors:** Jonathan H. Massey, Jun Li, David L. Stern, Patricia J. Wittkopp

## Abstract

Understanding the genetic basis of species differences is a major goal in evolutionary biology. Pigmentation divergence between *Drosophila* species often involves genetic changes in pigmentation candidate genes that pattern the body and wings, but it remains unclear how these changes affect pigmentation evolution in multiple body parts between the same diverging species. *Drosophila elegans* and *D. gunungcola* show pigmentation differences in the thorax, legs, and wings, with *D. elegans* exhibiting male-specific wing spots and *D. gunungcola* lacking wing spots with intensely dark thoraces and legs. Here, we performed QTL mapping to identify the genetic architecture of these differences. We find a large effect QTL on the X chromosome for all three body parts. QTL on Muller Element E were found for thorax pigmentation in both backcrosses but were only marginally significant in one backcross for the legs and wings. Consistent with this observation, we isolated the effects of the Muller Element E QTL by introgressing *D. gunungcola* alleles into a *D. elegans* genetic background and found that *D. gunungcola* alleles linked near the pigmentation candidate gene *ebony* caused intense darkening of the thorax, minimal darkening of legs, and minimal shrinking of wing spots. *D. elegans ebony* mutants showed changes in pigmentation consistent with Ebony having different effects on pigmentation in different tissues. Our results suggest that multiple genes have evolved differential effects on pigmentation levels in different body regions.

## Introduction

Pigmentation differences within and between species illustrate some of the most striking examples of phenotypic evolution in nature. In insects, pigmentation intensity in the body, legs, and wings varies widely, often distinguishing sexes, populations, and species (True, 2003; Wittkopp *et al*., 2003). The genetic and developmental processes determining pigment patterning are well understood, which has facilitated the use of insect pigmentation as a model system to investigate the genetic basis of phenotypic evolution (Wittkopp *et al*., 2003; Kopp, 2009). Multiple studies have revealed that the same genes have often evolved independently to cause pigmentation variation; that genetic variation within these genes often explains the majority of pigmentation differences; and that mutations affecting the expression of enzymes and transcription factors cause pigmentation to evolve (Massey and Wittkopp, 2016). These results support the hypothesis that the genetic architecture of phenotypic evolution is, at least to some degree, predictable (Stern and Orgogozo, 2008).

The insect pigmentation synthesis pathway (Figure 1) therefore provides a roadmap for predicting which genes likely contribute to pigmentation differences within and between insect species. Differences in how dopamine is metabolized in this pathway ultimately lead to combinations of black and brown melanins as well as yellow, tan, and colorless sclerotins that diffuse into the developing cuticle in sex-, population-, and species-specific ways (reviewed in True, 2003). The genes *yellow, tan*, and *ebony* in this pathway have repeatedly evolved to contribute to these differences. These include both within and between species differences in abdominal, thorax, and wing pigmentation (reviewed in Massey and Wittkopp, 2016). In each case, mutations in *cis*-regulatory DNA underlie these phenotypic changes, emphasizing the importance of gene regulatory evolution in generating animal pigmentation diversity.

**Figure 1.**
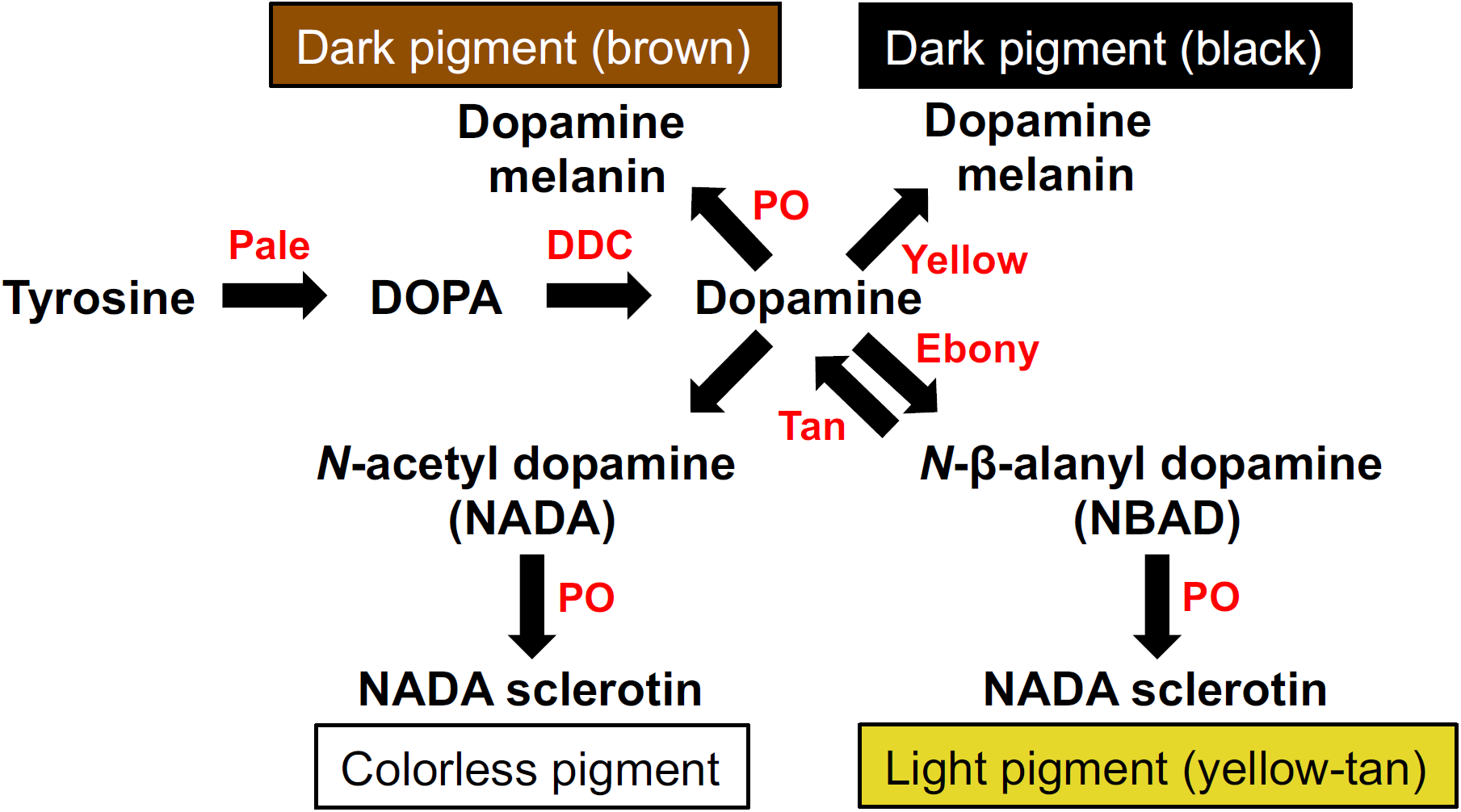
Insect sclerotization and pigmentation synthesis pathway. Pigmentation enzymes (shown in red) convert substrates (shown in black) into pigmentation precursors that polymerize into darkly colored melanins or lightly colored sclerotins. Synthesis of black melanin depends on Yellow function, whereas Ebony converts dopamine into lightly colored sclerotins. Tan catalyzes the reverse reaction of Ebony, converting NBAD molecules back into dopamine.

Most studies identifying the genetic basis of divergent pigmentation have focused on the divergence of one particular element of the pigment pattern within each species pair. It thus remains unclear how often the same genes and/or mutations are responsible for divergent pigmentation seen in multiple body parts. In some instances, closely related species show striking pigmentation differences across most body regions. Pigmentation divergence between *Drosophila americana* and *D. novamexicana*, for example, involves differences in thorax and abdominal pigmentation intensity that are partly explained by evolution of the *ebony* gene (Lamb *et al*., 2020). Divergence in pupal case coloration between *D. americana* and *D. virilis*, however, is caused by evolution at the *dopamine N-acetyltransferase* (*Dat*) gene (Ahmed-Braimah and Sweigart, 2015). In other instances, species differ not only in the pigmentation intensity across multiple body parts but also in pigmentation patterning. Pigmentation evolution in redheaded pine sawfly larvae, for example, involves changes in body coloration and spot patterning that is partially explained by overlapping quantitative trait loci (QTL) (Linnen *et al*., 2018).

Species differences between *D. elegans* and *D. gunungcola* involve dramatic changes in pigmentation intensity across multiple body parts, although body pigmentation is also polymorphic in *D. elegans* (Bock and Wheeler, 1972; Hirai and Kimura, 1997). Populations of *elegans* from Hong Kong and Indonesia have light brown thoraxes and legs, and males possess dark black spots at the tips of their wings (Hirai and Kimura, 1997; Figure 2). In Japan and Taiwan, populations have dark black thoraxes while still possessing male-specific wing spots (Hirai and Kimura, 1997). In *D. gunungcola*, pigmentation is less variable than in *D. elegans* (Sultana *et al*., 1999; Massey *et al*., 2020). Males and females have dark black thoraxes like the *D. elegans* Japanese and Taiwanese populations but also show dark black-pigmented legs, and males do not have wing spots (Yeh *et al*., 2006; Massey *et al*., 2020; Figure 2). Although these two species diverged 2-2.8 million years ago (Prud’Homme *et al*., 2006), they can produce fertile, female F_1_ hybrids in the laboratory (Yeh *et al*., 2006), allowing us to investigate the genetic basis of pigmentation evolution across multiple body regions in the same mapping population.

**Figure 2.**
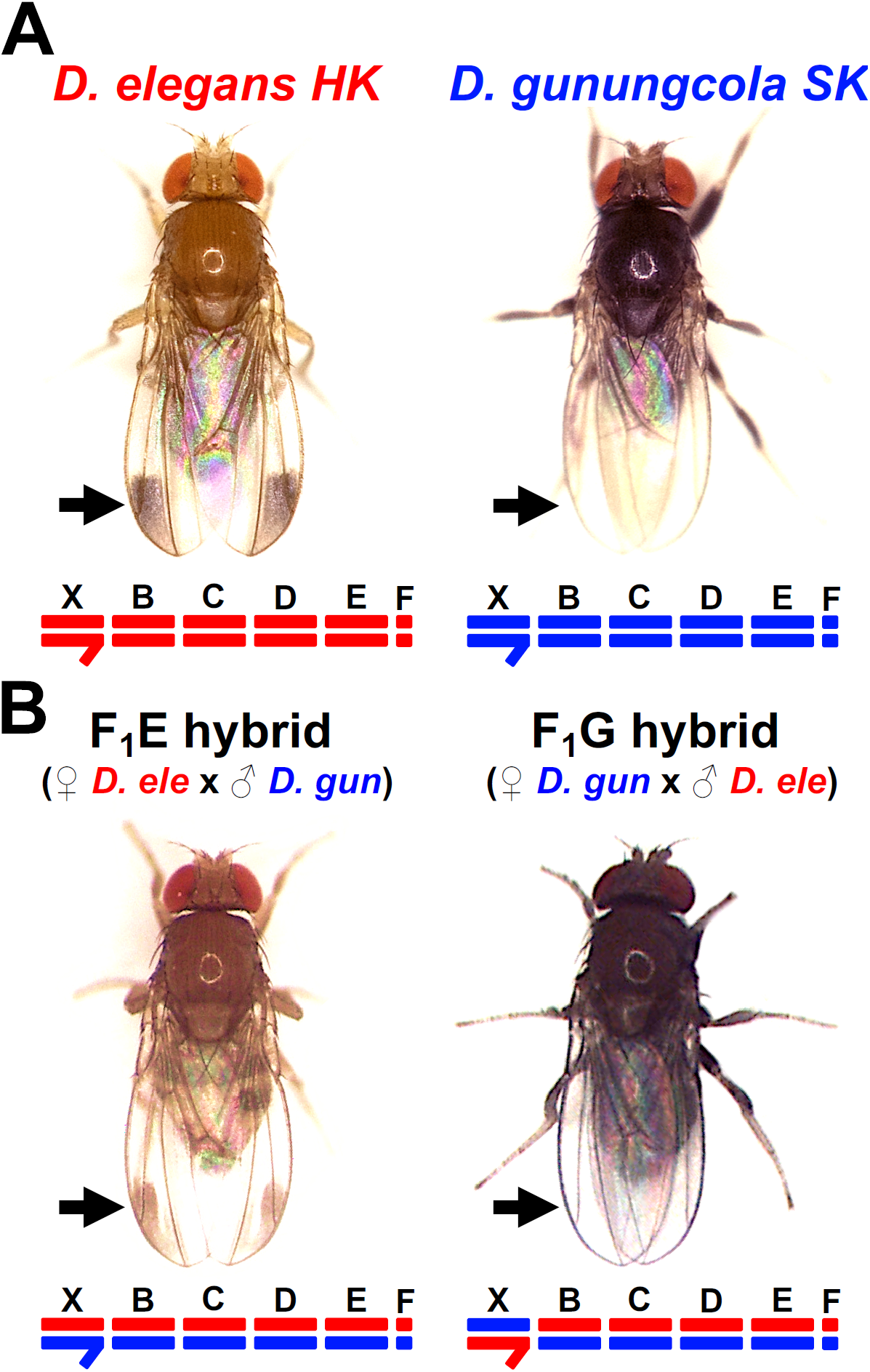
Differences in reciprocal F_1_male hybrids suggest X-linked sequence divergence in pigmentation evolution between *Drosophila elegans HK* and *D. gunungcola SK*. (A) *D. elegans HK (Hong Kong)* and *D. gunungcola SK (Sukarami)* show divergent thorax, leg, and wing pigmentation. (B) F_1_ male hybrids inheriting their X chromosome from *D. elegans HK* mothers (F_1_E, bottom left) possess wing spots and have thorax and leg pigmentation similar to *D. elegans HK*. F_1_ male hybrids inheriting their X chromosome from *D. gunungcola SK* mothers (F_1_G, bottom right) lack wing spots and have thorax and leg pigmentation similar to *D. gunungcola SK*.

Here, we use quantitative trait locus (QTL) mapping in two backcross populations between the light *D. elegans* morph from Hong Kong (HK) and *D. gunungcola* from Sukarami, Indonesia (SK) to identify gene regions associated with thorax, leg, and wing pigmentation divergence. For all three traits, we identified a major effect QTL on the X chromosome, where pigmentation candidate genes *omb* and *yellow* are located. We also find evidence for QTL linked to Muller Element E for all three traits, but these effects depend on genetic background. We attempted to isolate these effects by crossing *D. gunungcola* alleles into a *D. elegans* genetic background and by introgressing a ∼1.5 Mb region containing the *ebony* locus, which is found on Muller Element We find that homozygous *D. gunungcola* alleles linked to this introgression darkened thorax pigmentation intensity to *D. gunungcola* levels, but only minimally affected leg and wing pigmentation, consistent with the differences observed in QTL mapping data for the three traits. These data suggest that during the evolution of pigmentation across multiple body parts, epistasis influences the extent to which individual pigmentation loci affect pigmentation divergence in each tissue. In the case of *D. elegans* and *D. gunungcola*, evolution of at least two loci is necessary to cause species differences in leg and wing pigmentation, but evolution at a single locus is sufficient for divergence in thorax pigmentation.

## Materials and Methods

### Fly stocks

*Drosophila elegans HK (Hong Kong)* and *D. gunungcola SK (Sukarami)* species stocks were a gift from John True (Stony Brook University). Stock maintenance and food recipes are described in Massey *et al*. (2020). In brief, stocks were maintained at 23°C on a 12 h light-dark cycle. At the third instar larval stage of development, adults were transferred onto new food, and Fisherbrand filter paper (cat# 09-790-2A) was added to the larval vials to facilitate pupariation.

### Generating hybrid progeny

Male and female *D. elegans HK* flies will reproduce with male and female *D. gunungcola SK* in the laboratory to produce fertile F_1_ hybrid female and sterile F_1_ hybrid male offspring (Yeh *et al*., 2006). Creating these F_1_ hybrids in populations large enough for genetic analysis, however, is difficult. Detailed methods are described in Massey *et al*. (2020). In brief, both *D. elegans HK* and *D. gunungcola SK* species stocks were expanded to establish populations of more than 10,000 flies. Next, virgin males and females from each species were placed in heterospecific crosses (*D. elegans HK* males with *D. gunungcola SK* females and vice versa) in groups of ten males and females to generate fertile F_1_ hybrid female offspring. Dozens of these crosses were set to create ∼120 F_1_ hybrid female offspring. Since F_1_ hybrid males are sterile (Yeh *et al*., 2006; Yeh and True, 2014), the F_1_ hybrid females were used to generate two backcross populations for QTL mapping. Briefly, for the *D. elegans HK* backcross population, ∼60 F_1_ hybrid females were crossed in the same vial with ∼60 *D. elegans HK* males and transferred onto new food every two weeks for ∼2.5 months, resulting in 724 recombinant individuals. For the *D. gunungcola SK* backcross population, ∼60 F_1_ hybrid females were crossed in the same vial with ∼60 *D. gunungcola SK* males and transferred onto new food every two weeks for ∼2.5 months, resulting in 241 recombinant individuals.

### Pigmentation quantification

For thorax pigmentation QTL analysis, male recombinants from each backcross population were organized into three thorax pigmentation classes. The lightest recombinants showed thorax pigmentation intensities similar to *D. elegans HK* and were given a score of 0; recombinants with intermediate thorax pigmentation intensities were given a score of 1; and recombinants with dark thorax pigmentation intensities similar to *D. gunungcola SK* were given a score of 2 (Figure 3A). To quantify the effects of the Muller Element E introgression region on thorax pigmentation (Figure 4B,C), individuals were placed thorax-side up on Scotch double sided sticky tape on glass microscope slides (Fisherbrand) (cat# 12-550-15) and imaged at the same exposure using a Canon EOS Rebel T6 camera mounted to a Canon MP-E 65 mm macro lens equipped with a ring light. The images were then imported into ImageJ software (version 1.50i) (Wayne Rasband, National Institutes of Health, USA; http://rsbweb.nih.gov/ij/) and converted to 32 bit grayscale. Using the “straight line segment” tool, a line was drawn between the anterior scutellar bristles (Figure 4B white dashed box) to measure the mean grayscale value of the cuticle. Data shown in Figure 4C were inverted so that higher values indicated darker pigmentation. Raw data for Figure 3 and 4 are deposited on Dryad.

**Figure 3.**
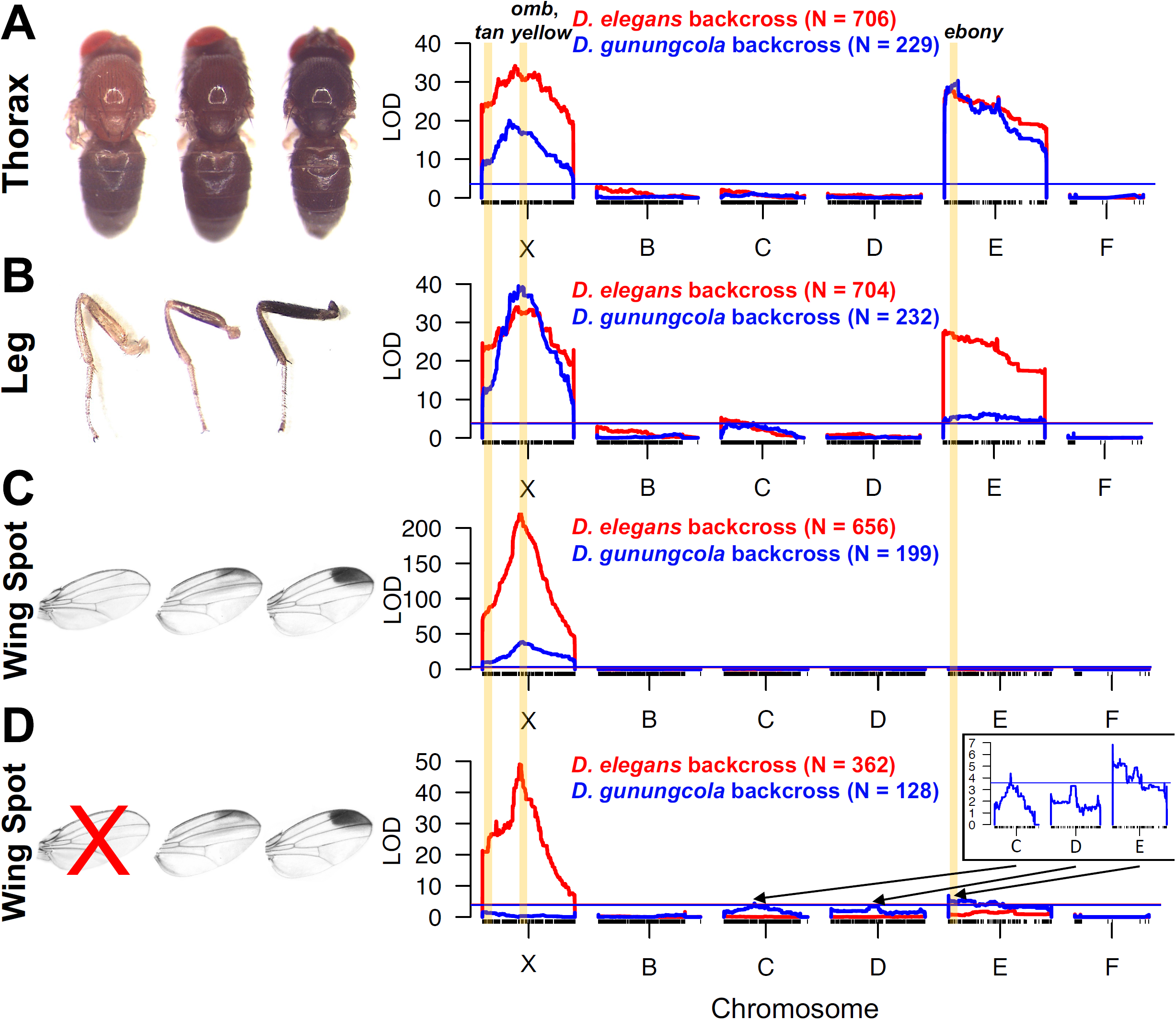
Quantitative trait locus (QTL) mapping of thorax, leg, and wing pigmentation divergence. (A) Thorax pigmentation was organized into three classes (light, intermediate, and dark). QTL map for thorax pigmentation is shown for *D. elegans HK* backcross (red) and *D. gunungcola SK* backcross (blue). (B) Leg pigmentation was organized into three classes (light, intermediate, and dark). QTL map for leg pigmentation is shown for *D. elegans HK* backcross (red) and *D. gunungcola SK* backcross (blue). (C) Wing spot pigmentation was quantified relative to wing size (see Methods). QTL map for wing spot pigmentation is shown for *D. elegans HK* backcross (red) and *D. gunungcola SK* backcross (blue) [Reprinted from Massey et al. (2020), Copyright (2020), with permission from John Wiley and Sons]. (D) Similar to (C), wing spot pigmentation was quantified relative to wing size, however, recombinants completely lacking wing spots were excluded (red “X”) from the QTL analysis. QTL map for wing spot size is shown for *D. elegans HK* backcross (red) and *D. gunungcola SK* backcross (blue) [Reprinted from Massey et al. (2020), Copyright (2020), with permission from John Wiley and Sons]. Transparent yellow bars indicate the chromosomal location of pigmentation candidate genes. LOD (logarithm of the odds) is indicated on the y-axis. The x-axis represents the physical map of Muller Elements X, B, C, D, E, and F based on the *D. elegans HK* assembled genome (see Methods). *D. elegans HK* and *D. gunungcola SK* have six chromosomes (Yeh *et al*. 2006; Yeh and True 2014) that correspond to *D. melanogaster* chromosomes as follows: X = X, B = 2L, C = 2R, D = 3L, E = 3R, and F = 4. Individual SNP markers are indicated with black tick marks along the x-axis. Horizontal red and blue lines mark P = 0.01 for the *D. elegans HK* and *D. gunungcola SK* backcross, respectively.

**Figure 4.**
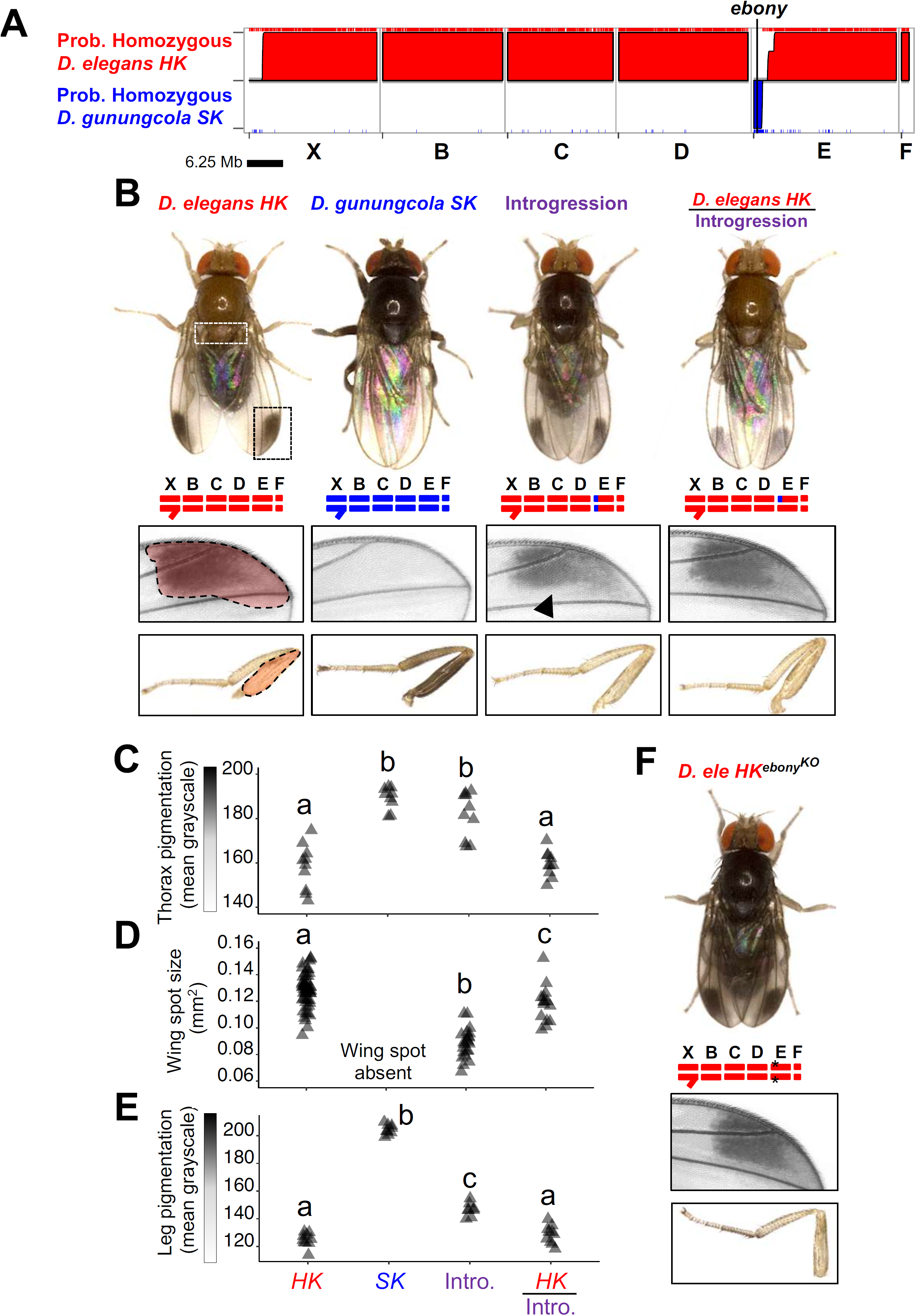
*D. gunungcola SK* alleles linked to *ebony* on Muller Element E have varied effects on thorax, leg, and wing pigmentation divergence. (A) Multiplexed Shotgun Genotyping (MSG) (Andolfatto et al., 2011) was used to estimate genome-wide ancestry assignments for a single introgression line generated by repeated backcrossing of dark *D. gunungcola SK* recombinant females with *D. elegans HK* males (see Methods). The posterior probability that a region is homozygous for *D. elegans HK* (red) or *D. gunungcola SK* (blue) ancestry is plotted along the y-axis. (B) Images highlighting pigmentation and chromosome differences between species and the Muller Element E introgression line. The white dashed box highlights the scutellar region of the thorax used to quantify thorax pigmentation in (C) (see Methods). Dashed red transparencies highlight regions of the wing and leg used to quantify wing spot and leg pigmentation in (D) and (E), respectively (see Methods). The black arrowhead emphasizes the wing spot region that has shrunk in the introgression line. (C) Quantification of thorax pigmentation differences between species and the Muller Element E introgression line (One-way ANOVA F_3,35_, = 33.9; P = 1.84 x 10^−10^; groups not sharing the same letter are significantly different based on post-hoc Tukey HSD at P < 0.00001). (D) Quantification of wing spot pigmentation differences between species and the Muller Element E introgression line (One-way ANOVA F_2,88_= 78.6; P < 2.0 x 10^−16^; groups not sharing the same letter are significantly different based on post-hoc Tukey HSD at P = 0.02) [Reprinted from Massey et al. (2020), Copyright (2020), with permission from John Wiley and Sons]. (E) Quantification of leg pigmentation differences between species and the Muller Element E introgression line. (One-way ANOVA F_3,33_ = 481; P < 2.0 x 10^−16^; groups not sharing the same letter are significantly different based on post-hoc Tukey HSD at P < 5 x 10^−7^). Gray triangles represent individual replicates for (C-E). (F) Images highlighting effects of knocking out the pigmentation candidate gene *ebony* in *D. elegans HK*.

For leg pigmentation QTL analysis, methods were identical to the thorax procedures above except recombinants were organized into light (0), intermediate (1), and dark (2) classes based on the pigmentation intensity of the medial side of their right hindleg femur (Figure 3B). This region of the leg was chosen because it contains few cuticular bristles that could obscure pigmentation. To quantify the effects of the Muller Element E introgression region on leg pigmentation (Figure 4B,E), the same methods as above were used except right medial hindleg images were captured. In ImageJ, the “polygon selections” tool was used to draw a polygon selection around the medial side of the right hindleg femur (Figure 4B red transparency selection) to measure the mean grayscale value of the cuticle. Data shown in Figure 4E were inverted so that higher values indicated darker pigmentation. Raw data for Figure 3 and 4 are deposited on Dryad.

For wing spot pigmentation QTL analysis, methods are described in Massey *et al*., (2020). Briefly, right wings from male recombinants from each backcross population were imaged, and spots were quantified in ImageJ using the “polygon selections” tool to quantify wing spot size relative to wing size (Figure 4B red transparency selection). The same procedure was used to quantify the effects of the Muller Element E introgression region (Figure 4D). Raw data for Figure 3 and 4 are deposited on Dryad.

### Library preparation, sequencing, and genome assembly

Detailed methods for preparing and sequencing the genomic DNA (gDNA) libraries for the *D. elegans HK* and *D. gunungcola SK* backcross populations and advanced introgression line were described in Massey *et al*., (2020). Briefly, gDNA was extracted from male recombinants by homogenizing individuals singly in each well of a 96-well plate (Corning, cat# 3879). Each recombinant gDNA sample was then barcoded with unique adaptors, pooled into a single multiplexed sequencing library, size selected, and sequenced in a single lane of Illumina HiSeq by the Janelia Quantitative Genomics Team. Methods for assembling the *D. elegans HK* and *D. gunungcola SK* genomes to facilitate marker generation were described in Massey *et al*. (2020).

### Marker generation with Multiplexed Shotgun Genotyping

Chromosome ancestry “genotypes” for the backcross progeny and introgression line were estimated with two Multiplexed Shotgun Genotyping (MSG) (Andolfatto et al., 2011) libraries, following methods described in Cande *et al*., (2012). Briefly, reads generated from the Illumina backcross sequencing library were mapped to the assembled *D. elegans HK* and *D. gunungcola SK* parental genomes to estimate chromosome ancestry for each backcross individual. We generated 3,425 and 3,121 markers for the *D. elegans HK* and *D. gunungcola SK* backcrosses, respectively, for QTL analysis [markers, phenotypes, and procedures for QTL mapping are deposited on Dryad (doi:10.5061/dryad.gb5mkkwm5)]. PDFs of chromosomal breakpoints for each recombinant are available here: https://deepblue.lib.umich.edu/data/concern/data_sets/j098zb17n?locale=en.

### QTL analysis

QTL analysis was performed using R/qtl (Broman and Sen, 2009) in R for Mac version 3.3.3. (R Core Team 2018). We imported ancestry data for both backcross populations into R/qtl using a custom script (https://github.com/dstern/read_cross_msg). This script directly imports the conditional probability estimates produced by the Hidden Markov Model (HMM) of MSG (described in detail in Andolfatto *et al*., 2011). We performed genome scans with a single QTL model using the “scanone” function of R/qtl and Haley-Knott regression (Haley and Knott, 1992) for thorax, leg, and wing pigmentation. For QTL mapping using the *D. elegans HK* backcross population, we excluded 18 and 20 individuals for thorax and leg pigmentation, respectively, because fly samples were either too poor to image or sequencing reads were too shallow to map. For the *D. gunungcola SK* backcross population, we excluded 12 and 9 individuals for thorax and leg pigmentation, respectively, for the same reasons. For wing pigmentation QTL analysis, we previously (Massey *et al*., 2020) reported on an ∼400 kb fine-mapped region on the X chromosome explaining the majority of variation for wing spot size. We also reported on QTL underlying variation in wing spot size with spotless recombinants removed from the analysis (Massey *et al*., 2020). Here, in Figure 3D, we show QTL underlying this variation to emphasize the role of Muller Element E in spot size divergence. Significance of QTL peaks at α = 0.01 was determined by performing 1000 permutations of the data.

### *Introgression of black body color alleles from* D. gunungcola SK *into* D. elegans HK

To isolate individual QTL underlying body color differences between *D. elegans HK* and *D. gunungcola SK*, F_1_ hybrid females and *D. elegans HK* backcross recombinants were generated using the methods described above. Next, dark black/brown female recombinants were repeatedly backcrossed with *D. elegans HK* males for four generations (BC3-BC6). Finally, we generated a single BC6 dark black homozygous introgression line that we then genotyped using MSG (Figure 4A,B)

*Creating an* ebony *null allele in* Drosophila elegans HK *via CRISPR-Cas9 genome editing* Using methods described in Bassett *et al*. 2013, we *in vitro* transcribed (MEGAscript T7 Transcription Kit, Invitrogen) three single guide RNAs (sgRNAs) (Supplemental File S1) with target sequences designed based on conserved sites between *D. elegans HK* and *D. gunungcola SK* in exon 2 of the *ebony* gene. After transcription, sgRNAs were purified using an RNA Clean and Concentrator 5 kit (Zymo Research), eluted with nuclease-free water, and quantified using a Qubit RNA BR Assay Kit (Thermo Fisher Scientific). Next, mature (>2 weeks old) *D. elegans HK* males and females were transferred to 60 mm embryo lay cages (GENESEE Part Number: 59-100) on top of 60 mm grape plates (3% agar + 25% grape juice + 0.3% sucrose) at high densities (>300 flies) after brief CO_2_ anesthesia. After CO_2_ anesthesia, mature, mated *D. elegans HK* females will often dispel an embryo from their abdomen where it sticks briefly to the anus. Tapping the grape plate + embryo cage down on a hard surface 10 times causes the embryos to stick to the grape agar. Flies were then transferred back into food vials and embryos were lined up on glass cover slips taped to a glass microscope slide. For CRISPR/Cas9 injections into *D. elegans HK* embryos, Cas9 protein (PNA Bio #CP01), phenol red, and all three sgRNAs were mixed together at 400 ng/µl, 0.05%, and 100 ng/µl final injection concentrations, respectively. All CRISPR injections were performed in-house, using previously described methods (Miller *et al*, 2002). Finally, we screened for germline mutants based on body pigmentation and confirmed loss of Ebony protein by western blot (Supplemental Figure S1). To the best of our knowledge, these are the first gene editing experiments to succeed in *D. elegans*. We attempted the same experiments in *D. gunungcola SK*, but failed to recover any mutants.

### Western blotting

Western blot methods were followed similar to Wittkopp *et al*. (2002). In brief, for each replicate per genotype, four newly eclosed (within 60 min) male flies were homogenized in 100 µl of 125 mM Tris pH 6.8, 6% SDS and centrifuged for 15 min. The supernatant was then transferred to a new protein low-bind Eppendorf tube with an equal amount of 2X Laemmli sample buffer [4% SDS, 20% glycerol, 120 mM Tris-Cl (pH 6.8)], boiled for 10 min, and stored at -80°C. Before gel electrophoresis, samples were thawed at room temperature for 30 min, and 20 µl of each sample was loaded into individual wells of an Invitrogen NuPAGE 4-12% Bis-Tris Gel. The gel was run at 175 V for 60 min, washed, and transferred to an Invitrogen iBlot 2 PVDF Mini Stack Kit. The mini stack was run on an iBlot 2 (ThermoFisher, Catalog Number: IB21001) to perform western blotting transfer, and blocked using an Invitrogen Western Breeze anti-rabbit kit (Catalog Number: WB7106). After two washes with diH_2_O, samples were incubated in 1:400 rabbit anti-Ebony (Wittkopp *et al*., 2002) overnight at 4°C. Finally, samples were washed using the Western Breeze wash solution, incubated in secondary antibody solution alk-phos. (Catalog Number: WP20007) with conjugated anti-rabbit for 30 min, washed for 2 min with diH_2_O, and prepared for chromogenic detection.

### Statistics

Statistical tests were performed in R for Mac version 3.3.3 (R Core Team 2018). ANOVAs were performed with post hoc Tukey HSD for pairwise comparisons adjusted for multiple comparisons. See “QTL analysis” methods for statistical tests used during QTL mapping.

## Results

### X-linked divergence contributed to pigmentation divergence of all three traits

We examined the genetic basis for divergence of thorax, leg, and wing pigmentation between *D. elegans HK* and *D. gunungcola SK* (Figure 2A). First, to assess the contribution of the X chromosome in pigmentation divergence, we compared F_1_ hybrid males from reciprocal crosses between *D. elegans HK* and *D. gunungcola SK*, which inherited their X chromosome from either their *D. elegans HK* (F_1_E) or *D. gunungcola SK* (F_1_G) mother, respectively. We observed striking pigmentation differences in the thorax, legs, and wings between the hybrids (Figure 2B). F_1_ hybrid males inheriting their X chromosome from *D. elegans HK* mothers possessed wing spots (quantified in Massey *et al*., 2020) and had thorax and leg pigmentation similar to *D. elegans HK* (Figure 2A,B). In contrast, F_1_ hybrids from *D. gunungcola SK* mothers lacked wing spots and had thorax and leg pigmentation similar to *D. gunungcola SK* (Figure 2A,B). These observations suggest that X-linked loci contributed to pigmentation divergence in multiple body parts.

### QTL on the X chromosome underlie pigmentation divergence

To assess the genome-wide distribution of genes contributing to pigmentation divergence for each body part, we performed quantitative trait locus (QTL) mapping. We generated two backcross populations by crossing F_1_ hybrid females to males from each parental species and quantified pigmentation variation for thorax, leg, and wing spots (see Methods) (Figure 3). For each trait in both backcrosses, we identified major effect QTL on the X chromosome (Figure 3; Table 1). The X-linked QTL peaks for thorax and leg pigmentation appear broader than the wing spot pigmentation QTL (Figure 3), which is also reflected in their support intervals (Table 1). For the wing spot, this QTL was previously refined to a ∼400 kb region containing the pigmentation candidate gene *optomotor-blind* (*omb*) (Massey *et al*., 2020; Table 1), consistent with the single, narrow peak for wing spot variation (Figure 3C). The broader QTL peaks for variation in thorax and leg pigmentation may result from multiple X-linked genes affecting these traits.

**Table 1.**
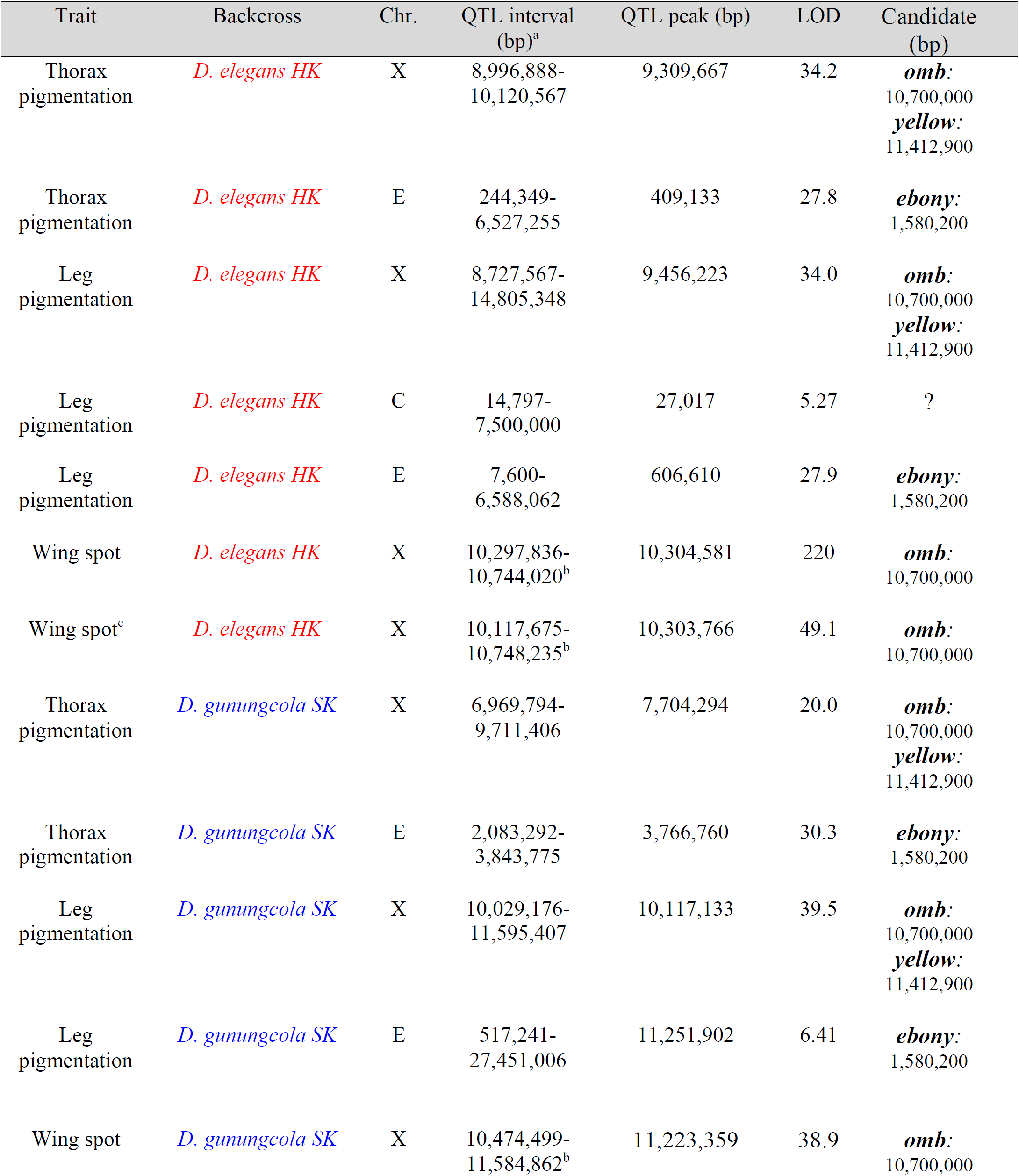
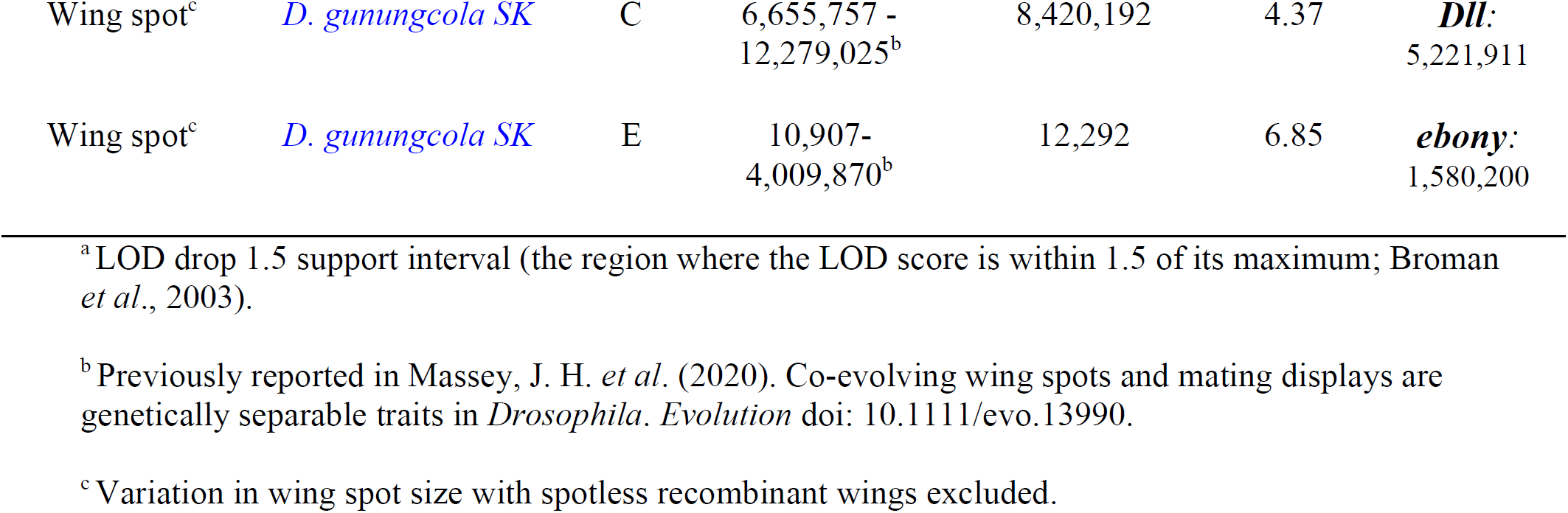
QTLs detected for thorax, leg, and wing pigmentation divergence.

### QTL on Muller Element E underlie pigmentation divergence

We detected major effect QTL for thorax and leg, but not wing, pigmentation on Muller Element E (Figure 3A,B), with the strongest effect linked to the left end of the chromosome (Table 1). For these traits, the QTL peaks cover the entire Muller element, which might indicate low recombination rate or the presence of multiple genes on Muller Element E contributing to thorax and leg pigmentation divergence. Nonetheless, this result indicates that at least one gene on Muller Element E affects thorax and leg pigmentation divergence differently than wing spot pigmentation. Prior work has also shown, however, that autosomal QTL on Muller Elements C and E affect the size of wing spots even though the X chromosome is sufficient to control their presence or absence (Massey et al, 2020; Figure 3D).

To investigate further whether autosomal variation might affect wing spot divergence, and whether this variation might be hidden by a large epistatic effect of the X chromosome, we repeated QTL analysis on a subset of backcross recombinants that displayed wing spots of any size and excluded offspring that completely lacked spots (Figure 3D). This analysis revealed small effect QTL linked to Muller Elements C and E that affect wing spot size (Figure 3D). Like thorax and leg pigmentation QTL, the largest effect was linked to the left end of Muller Element E.

### *ebony* is a candidate gene for the Muller Element E QTL

The pigmentation gene *ebony* is located near the left end of Muller Element E, where we observed linkage to thorax, leg, and wing spot size pigmentation differences in the backcross populations, making it an excellent candidate for the locus harboring the genetic variant(s) affecting these traits (Figure 3). To test this hypothesis, we attempted to create stable introgression lines by repeatedly backcrossing *D. gunungcola SK* alleles into a *D. elegans HK* genetic background. For each backcross, we selected for dark black female recombinants to cross with virgin male *D. elegans HK* for six generations (see Methods). Our goal was to isolate multiple, independent introgression lines to fine map QTL involved in pigmentation divergence; however, the majority of introgressions failed to propagate. We were not able to maintain any introgression lines containing *D. gunungcola SK* alleles linked to the X chromosome in a *D. elegans HK* genetic background (Massey *et al*., 2020), suggesting that divergence on the X chromosome also contributed to hybrid incompatibilities. We did, however, recover a single introgression line with a dark black thorax that resembled *D. gunungcola SK* (Figure 4A,B). Sequencing this line revealed that it was homozygous for an approximately 1.5 Mb region of the left end of *D. gunungcola SK* Muller Element E loci that included the *ebony* gene (Figure 4A). These data provide independent confirmation that variation in genes on the left end of Muller Element E contribute to thorax pigmentation divergence and support the hypothesis that *ebony* contributes to thorax pigmentation evolution.

We next quantified pigmentation variation of the Muller Element E introgression line to determine whether genetic variation in this region is sufficient to explain the Muller Element E QTL effects. We found that this introgression completely recapitulated the species difference in thorax pigmentation, but it had only weak effects on leg pigmentation and wing spot size (Figure 4C-E). This result allowed us to partially separate the genetic effects on Muller Element E, further implicating *ebony* as a candidate gene with a large effect on thorax pigmentation divergence and a small effect on leg and wing spot pigmentation.

To determine whether *ebony* contributes to thorax pigmentation divergence, we attempted to knock out the *ebony* gene in both species using CRISPR-Cas9 genome editing to perform a reciprocal hemizygosisty test (Stern, 2014). Unfortunately, we were able to generate only a single *ebony* knockout line in *D. elegans HK* (Figure 4F; Supplemental Figure S1). We found that the *D. elegans HK ebony* line displayed a dark black thorax (Figure 4F) from loss of Ebony activity, similar to *ebony* mutant effects in *D. melanogaster* (Wittkopp *et al*., 2002). This result further supports the hypothesis that lower Ebony activity and/or expression in *D. gunungcola SK* underlies its dark thorax color. Interestingly, loss of Ebony activity did not darken leg pigmentation or expand the size of wing spots in *D. elegans HK* (Figure 4F). One explanation for this observation is that *ebony* is not normally expressed in these tissues in *D. elegans HK*, so loss of Ebony function is not reflected phenotypically in these regions. Alternatively, because pigmentation reflects the balance of expression between *ebony, tan*, and *yellow* (Wittkopp *et al*., 2002; Wittkopp *et al*., 2009), differences in expression of these other genes might explain why we did not observe the accumulation of dark black melanins in these tissues of *D. elegans* HK *ebony* mutants (Figure 1). This observation could explain why genetic variation linked near the *ebony* locus seems to affect thorax pigmentation divergence more than leg and wing pigmentation divergence.

## Discussion

We found that pigmentation divergence in the thorax, legs, and wings between *D. elegans HK* and *D. gunungcola SK* is primarily influenced by genes on the X chromosome and Muller Element E. Despite the fact that QTL regions overlap for these three body regions, several observations suggest that pigmentation divergence results from evolution of multiple genes that act in different ways on the three body regions examined. First, the X-linked QTLs that influence thorax and leg pigmentation are broader than the QTL for wing spots. Second, QTL on Muller Element E have a strong effect on thorax and leg pigmentation but minimal effect on wing spots. Third, a 1.5 Mbp introgression of Muller Element E had a strong effect on thorax pigmentation but minimal effects on leg pigmentation and wing spots. Thus, there is evidence for independent genetic variation influencing pigmentation in each of these three body regions.

Evidence from previous studies is similar to our results. In *D. melanogaster*, for example, genetic variants in the *tan* gene affect pigmentation in multiple abdominal segments, but genetic variants in other genes affect each segment independently (Dembeck *et al*., 2015). Similarly, a single, large effect QTL changes bristle number across multiple body regions in *D. melanogaster*, but QTL on different chromosomes have independent effects on bristle number in the thorax versus the abdomen (Mackay, 1995). Genetic variation in individual genes, therefore, can have pleiotropic effects on phenotypic differences in multiple body parts, whereas genetic variation in less pleiotropic genes can have minimal effects in specific body parts.

Multicellular organisms develop distinct body parts through the action of gene regulatory networks (Davidson, 2010). Differences in which (and how) genes are used during phenotypic evolution likely reflect the activity of these networks (Stern and Orgogozo, 2008). If a gene is not expressed in a given tissue, for example, then genetic variation at this gene will not be reflected phenotypically. In our study, QTL on the X chromosome influenced pigmentation differences in all three body parts (Figure 3), but QTL linked to the left end of Muller Element E mainly affected thorax pigmentation (Figure 4). We predict, therefore, that species differences in the spatial expression pattern of genes with distinct functions in the insect sclerotization and pigmentation synthesis pathway (Figure 1) underlie these results.

In *Drosophila*, pigmentation coloring and patterning is determined by the action of multiple enzymes that are regulated by multiple transcription factors (reviewed in Massey and Wittkopp, 2016). The enzymes Yellow and Ebony both require dopamine to synthesize dark black or light yellow pigments, respectively, and transcription factors determine where these enzymes are expressed (Figure 1; Massey and Wittkopp, 2016). Mutations that disrupt Yellow activity cause flies to develop light yellow pigments, because Ebony converts excess dopamine into lightly colored sclerotins. Reciprocally, mutations that disrupt Ebony activity cause flies to develop dark black pigments, because Yellow converts excess dopamine into dark black melanins (Figure 1). The phenotypic effect of mutations in one enzyme, therefore, require the reciprocal activity of the other (Wittkopp *et al*., 2002). When we introgressed *D. gunungcola SK* Muller Element E alleles containing the *ebony* gene into a *D. elegans HK* genetic background, we observed darkening of the thorax but not the leg or wing (Figure 4B). This likely resulted from the combined effects of low *D. gunungcola SK ebony* expression with high *D. elegans HK yellow* expression. We hypothesize, then, that the *D. elegans HK yellow* allele is not normally expressed in the legs or in regions outside the wing spot. Instead, the *D. gunungcola SK yellow* allele likely evolved an expanded expression pattern in the legs and a restricted expression pattern in the wings (Prud’homme *et al*., 2006). Dissecting these mechanisms in detail will shed light on how the evolution of gene regulatory networks shape phenotypic divergence and vice versa.

## Acknowledgements

We thank members of the Wittkopp, Stern, and Rebeiz labs for helpful discussions. For fly strains, we thank J. True (Stony Brook University). For guidance with CRISPR/Cas9 genome editing and embryo injections, we thank Abigail Lamb. For helpful advice on creating F_1_ hybrids, we thank Shu-Dan Yeh (National Central University). Funding was provided by University of Michigan, Department of Ecology and Evolutionary Biology, Peter Olaus Okkelberg Research Award, National Institutes of Health (NIH) training grant T32GM007544, and Howard Hughes Medical Institute Janelia Graduate Research Fellowship to JHM; NIH R01 GM089736 and 1R35GM118073 to PJW.

## Data Archiving

All supporting data can be accessed at University of Michigan Deep Blue (https://deepblue.lib.umich.edu/data/concern/data_sets/j098zb17n?locale=en) and Dryad.

## Figure Legends

**Supplementary Figure S1.**
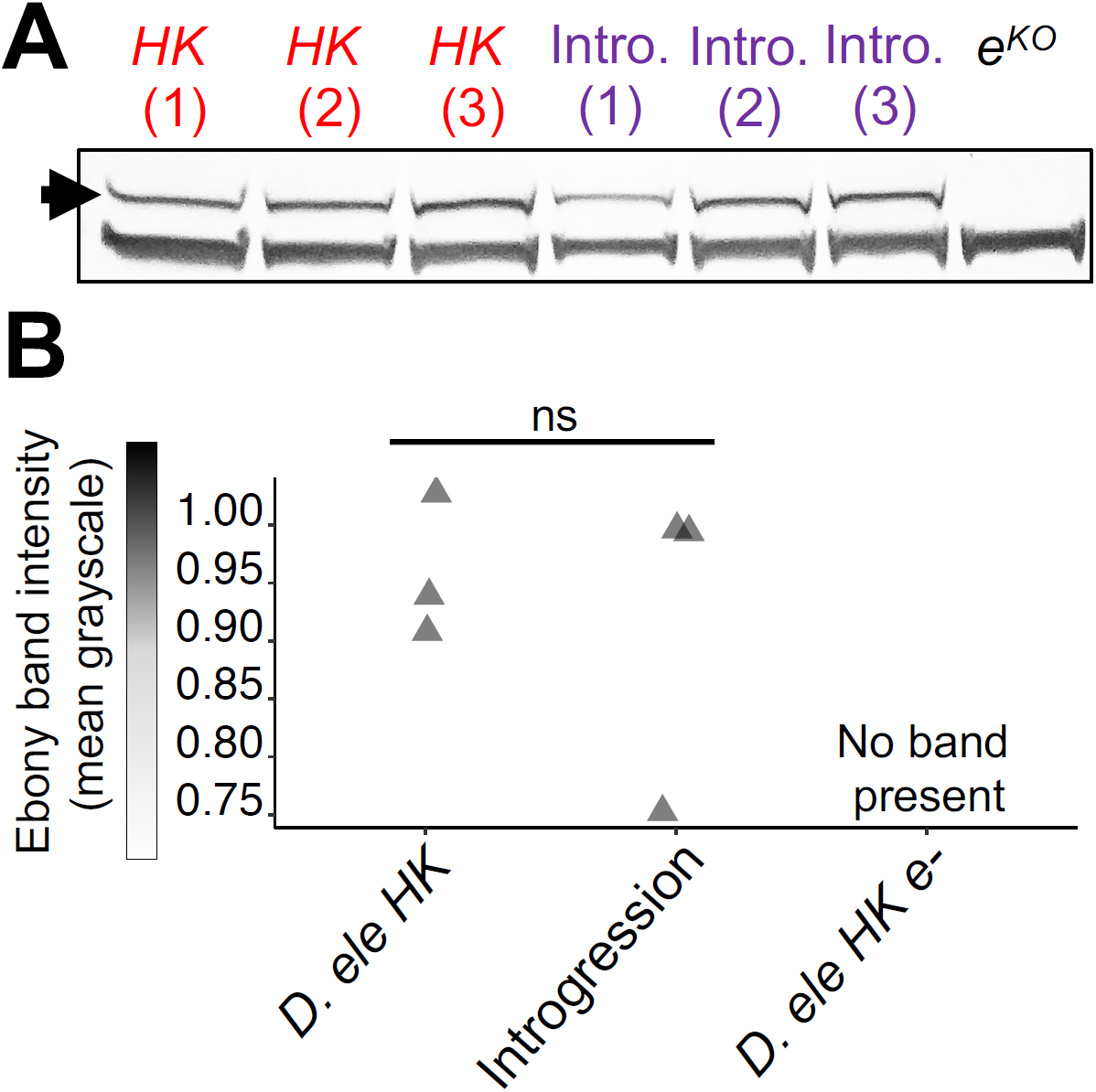
Quantification of Ebony band intensity in western blot. (A) Western blot using anti-Ebony antibody to detect Ebony protein levels (black arrowhead) in newly eclosed *D. elegans HK*, the Muller Element E introgression line, and the *D. elegans HK ebony* knockout line. Each number denotes a unique biological replicate involving four homogenized flies. A second band just below the ∼94 kDa Ebony band appeared in both the wild-type and *ebony* knockout extracts (as observed in Wittkopp *et al*., 2002). (B) Using this secondary band as a loading control, we quantified Ebony protein levels for each genotype and observed no statistically significant differences. The western blot image was imported into ImageJ software (version 1.50i) (Wayne Rasband, National Institutes of Health, USA; http://rsbweb.nih.gov/ij/) and converted to 32 bit grayscale. Using the “straight line segment” tool, the mean grayscale value of each band was quantified and inverted so that higher values indicated darker band intensity (Student’s t-test; t = -0.495; df = 2.75; P = 0.646; two-tailed). Gray triangles represent individual replicates, ns = not significant.

## References

Ahmed-Braimah, Y. H., & Sweigart, A. L. (2015). A single gene causes an interspecific difference in pigmentation in Drosophila. Genetics, 200(1), 331–342.

Andolfatto, P., Davison, D., Erezyilmaz, D., Hu, T. T., Mast, J., Sunayama-Morita, T., & Stern, D. L. (2011). Multiplexed shotgun genotyping for rapid and efficient genetic mapping. Genome research, 21(4), 610–617.

Bock, I. R., and Wheeler, M. R. (1972) The Drosophila melanogaster species group. University of Texas Publication 7213, 1–102.

Broman, K. W., & Sen, S. (2009). A Guide to QTL Mapping with R/qtl (Vol. 46). New York: Springer.

Davidson, E. H. (2010). The regulatory genome: gene regulatory networks in development and evolution. Elsevier.

Dembeck, L. M., Huang, W., Magwire, M. M., Lawrence, F., Lyman, R. F., & Mackay, T. F. (2015). Genetic architecture of abdominal pigmentation in Drosophila melanogaster. PLoS Genet, 11(5), e1005163.

Geyer, P. K., & Corces, V. G. (1987). Separate regulatory elements are responsible for the complex pattern of tissue-specific and developmental transcription of the yellow locus in Drosophila melanogaster. Genes & Development, 1(9), 996–1004.

Haley, C. S., and S. A. Knott. 1992. A simple regression method for mapping quantitative trait loci in line crosses using flanking markers. Heredity, 69:315–324.

Hirai, Y., and Kimura, M. T. (1997) Incipient reproductive isolation between two morphs of Drosophila elegans (Diptera: Drosophilidae). Biol. J. Linn. Soc., 61, 501–513.

Jeong, S., Rebeiz, M., Andolfatto, P., Werner, T., True, J., & Carroll, S. B. (2008). The evolution of gene regulation underlies a morphological difference between two Drosophila sister species. Cell, 132(5), 783–793.

Kopp, A. (2009). Metamodels and phylogenetic replication: a systematic approach to the evolution of developmental pathways. Evolution, 63(11), 2771–2789.

Lamb, A. M., Wang, Z., Simmer, P., Chung, H., Wittkopp, P. J. 2020. ebony affects pigmentation divergence and cuticular hydrocarbons in Drosophila americana and D. novamexicana. Front. Ecol. Evol. doi:10.3389/fevo.2020.00184

Linnen, C. R., O’Quin, C. T., Shackleford, T., Sears, C. R., & Lindstedt, C. (2018). Genetic basis of body color and spotting pattern in redheaded pine sawfly larvae (Neodiprion lecontei). Genetics, 209(1), 291–305.

Liu, Y., Ramos-Womack, M., Han, C., Reilly, P., Brackett, K. L., Rogers, W., … & Rebeiz, M. (2019). Changes throughout a genetic network mask the contribution of Hox gene evolution. Current Biology, 29(13), 2157–2166.

Mackay, T. F. (1995). The genetic basis of quantitative variation: numbers of sensory bristles of Drosophila melanogaster as a model system. Trends in Genetics, 11(12), 464–470.

Massey, J. H., & Wittkopp, P. J. (2016). The genetic basis of pigmentation differences within and between Drosophila species. In Current topics in developmental biology (Vol. 119, pp. 27–61). Academic Press.

Massey, J. H., Rice, G. R., Firdaus, A., Chen, C. Y., Yeh, S. D., Stern, D. L., Wittkopp, P. J. (2020). Co-evolving wing spots and mating displays are genetically separable traits in Drosophila. Evolution, DOI: doi:10.1111/evo.13990.

Miller, D. F., Holtzman, S. L., & Kaufman, T. C. (2002). Customized microinjection glass capillary needles for P-element transformations in Drosophila melanogaster. Biotechniques, 33(2), 366–375.

Prud’Homme, B., Gompel, N., Rokas, A., Kassner, V. A., Williams, T. M., Yeh, S. D., … & Carroll, S. B. (2006). Repeated morphological evolution through cis-regulatory changes in a pleiotropic gene. Nature, 440(7087), 1050–1053.

R Core Team. 2018. R: a language and environment for statistical computing. Available via http://www.r-project.org/

Rebeiz, M., Pool, J. E., Kassner, V. A., Aquadro, C. F., & Carroll, S. B. (2009). Stepwise modification of a modular enhancer underlies adaptation in a Drosophila population. Science, 326(5960), 1663–1667.

Stern, D. L. (2014). Identification of loci that cause phenotypic variation in diverse species with the reciprocal hemizygosity test. Trends in Genetics, 30(12), 547–554.

Stern, D. L., & Orgogozo, V. (2008). The loci of evolution: how predictable is genetic evolution?. Evolution, 62(9), 2155–2177.

Sultana, F., Kimura, M. T., & Toda, M. J. (1999). Anthophilic Drosophila of the elegans species-subgroup from Indonesia, with description of a new species (Diptera: Drosophilidae). Entomological Science, 2, 121–126.

True, J. R. (2003). Insect melanism: the molecules matter. Trends in Ecology & Evolution, 18(12), 640–647.

Wittkopp, P. J., True, J. R., & Carroll, S. B. (2002). Reciprocal functions of the Drosophila Yellow and Ebony proteins in the development and evolution of pigment patterns. Development, 129(8), 1849–1858.

Wittkopp, P. J., Carroll, S. B., & Kopp, A. (2003). Evolution in black and white: genetic control of pigment patterns in Drosophila. Trends in Genetics, 19(9), 495–504.

Wittkopp, P. J., Stewart, E. E., Arnold, L. L., Neidert, A. H., Haerum, B. K., Thompson, E. M., … & Shefner, L. (2009). Intraspecific polymorphism to interspecific divergence: genetics of pigmentation in Drosophila. Science, 326(5952), 540–544.

Yeh, S. D., Liou, S. R., & True, J. R. (2006). Genetics of divergence in male wing pigmentation and courtship behavior between Drosophila elegans and D. gunungcola. Heredity, 96(5), 383–395.

